# Comparison of the CDC2-like kinase family across eukaryotes highlights the functional conservation of these unique biological thermometers

**DOI:** 10.1101/2024.06.21.599975

**Authors:** Rachel A. Ogle, Jacob K. Netherton, Benjamin R. Robinson, Florian Heyd, Xu Dong Zhang, Mark A. Baker

**Affiliations:** Faculty of Science and Faculty of Health and Medicine, University of Newcastle, Callaghan NSW 2308, Australia; Institut für Chemie und Biochemie, RNA Biochemie, Freie Universität Berlin, Berlin, Germany

## Abstract

The family of CDC2-like kinases (CLKs) play a crucial role in regulating alternative splicing (AS), a process fundamental to eukaryotic gene expression and adaptation. Of particular interest, these enzymes exhibit unique responsiveness to minor temperature shifts, enabling them to modulate AS accordingly. Dysregulated CLK expression is linked to a wide variety of human diseases, establishing them as promising therapeutic targets. Despite the importance of CLKs, limited research has explored the genetic and functional diversification of this gene family. This report investigates the evolutionary origins, diversification, and functional implications of CLKs across major eukaryotic lineages through phylogenetic and structural comparisons. Our data demonstrate these kinases are prevalent throughout eukaryotes, with the original gene (which shares orthology to human CLK2), dating back to the Last Eukaryotic Common Ancestor. We identified three key duplication events in vertebrates, highlighting how this gene family has expanded and diversified in complex metazoans. Despite two instances of CLK paralog loss in vertebrate lineages, CLKs remain prevalent throughout metazoans, suggesting they are essential for complex eukaryotic life. Structural comparisons across diverse eukaryotes demonstrate kinase domain conservation, which is in line with their maintained function in AS regulation. While their N-terminal regions vary significantly in amino acid sequence, the function of this domain to regulate phosphorylation of AS factors is conserved, albeit in a species-specific manner. CLKs exhibit unique thermo-sensitive properties across diverse species, challenging conventional enzymatic behaviour. This temperature regulation, mediated by their kinase activation segment, is characterised by increased activity at lower physiological temperatures. The conservation of this structure, and a thermo-sensitive amino acid motif within it, suggests this was an ancient adaptation for responding to environmental cues. Species-specific temperature profiles highlight the adaptive evolution of CLKs, enabling organisms to thrive in diverse environmental conditions including extreme temperatures. Our analysis expands the understanding of CLK biology across diverse eukaryotes and connects insights from model organisms to human biology.

## Introduction

The family of Cdc2-like kinases (CLKs) possess dual-specificity, enabling them to phosphorylate serine/threonine, and tyrosine residues. Despite sharing structural similarities with Cdc2, CLKs have distinct functions. Throughout eukaryotes, CLK homologs are also termed LAMMER kinases due to the presence of the conserved “EHLAMMERILG” motif, and have been studied in human^1,2^, mouse^3^, fruit fly^4^, roundworm^5^, turtle^6^, alligator^6^, frog^7,8^, plants^9,10^, and yeast^11,12^. The number of genes in this family increases with organism complexity, with mammals possessing a total of four paralogs (CLK1, CLK2, CLK3 and CLK4). However, limited research has explored the genetic and functional diversification of this gene family. While some partial phylogenetic analyses of CLKs have been performed elsewhere^10,12–14^, to date, no study has demonstrated a complete eukaryotic timeline, which could offer valuable insights in predicting functional roles and diversification across species^15^. Similarly, whilst crystal structure comparisons of human CLK1-4 have been determined^1,2,16,17^, there exists a gap in evaluating the structural conservation of CLK homologs between different eukaryotes.

An important and well-known function of the CLK family, conserved from mammals to plants, is their role in regulating both constitutive and alternative pre-mRNA splicing through the phosphorylation of serine/arginine-rich proteins (SR proteins)^2,10,18,19^. Alternative splicing (AS) is a fundamental process in multicellular eukaryotes, facilitating the production of multiple protein isoforms from a single gene and contributing to their ability to adapt and diversify^20^. The importance of AS is further highlighted by the fact that mutations affecting splice site recognition account for about 15% of all hereditary disease-causing mutations in humans^21^. Inhibition of CLK causes widespread changes to global AS^6,10^. Hence, it is not surprising that aberrant CLK expression in humans has been extensively linked to various diseases, including cancer, muscular dystrophy, Alzheimer’s, osteoarthritis, and viral replication (reviewed in detail elsewhere ^1,2,22^). For this reason, CLKs are being rapidly established as effective targets for therapeutic intervention, highlighting the importance of understanding their functional biology.

Organisms continuously utilise AS to rapidly shift their intercellular processes in response to environmental cues. In the context of CLKs, a notable function is their ability to regulate AS in response to small changes in temperature. This newly discovered feature is especially unique, such that their activity is upregulated by just a 1°C decrease in physiological temperature, which is in stark contrast to typical kinase thermodynamics. This phenomenon has been observed in homologs across diverse eukaryotes, including human, mouse, fruit fly, turtle, alligator, plant, and thermophilic algae^6,10,23^, suggesting this mechanism is likely of ancient eukaryotic origin. This unique adaptation allows CLKs to promptly alter AS in a temperature-dependant manner by modulating SR protein phosphorylation. Currently, this function has potential links to the regulation of important biological processes, including circadian rhythm cycles in mammals, reptilian temperature-dependent sex determination, and plant thermomorphogenesis^6,10,24^. Considering that all eukaryotic organisms experience temperature fluctuations, this positions CLKs as potential regulators of an immense range of biological processes. Remarkably, the temperature profile at which CLK homologs are regulated has adapted to the specific physiological environment of each host organism. For example, in mammals, activity of CLK1/4 is typically regulated between 33-38°C. However, in organisms that inhabit more extreme conditions, such as the thermophilic red algae *Cyanidioschyzon merolae*, temperature-regulation of its CLK homolog “LIK” occurs between 48-56°C^23^. Despite variations in their activity profile, a key aspect to their functionality is that all CLKs examined so far exhibit increased activity below an organism’s physiological temperature.

This paper aims to address gaps in our knowledge of the evolution and diversification of the CLK family of kinases. Our approaches include: (i) a phylogenetic analysis demonstrating gene expansion and gene loss, (ii) structural and functional comparisons of their kinase domain and N-terminal to investigate their conservation, and (iii) an investigation into how their unique temperature regulatory mechanisms have evolved and diversified. Although the importance of CLKs in the context of humans is rapidly expanding due to their association with various diseases, there is a substantial body of research in other model species that could shed light on their function. Thus, this work offers insights into the functional evolution of the CLK family, serving as a resource for researchers interested in studying these kinases across eukaryotic organisms.

## Results and Discussion

### (1) Phylogenetic history of CLKs

#### Origins of the CLK family

The CLK family is suggested to have originated during eukaryogenesis, with a single CLK2-related gene being present in the Last Eukaryotic Common Ancestor (LECA)^25^. To understand their diversification, a phylogenetic analysis using 64 CLK homologs across 28 eukaryotic organisms was undertaken (Figure 1). The data show that the ancestral kinase preceding the expansion of this family shares orthology with CLK2, consistent with findings from the Eukaryotic Kinome Tree^25^. Moreover, we found no evidence of homologous proteins within prokaryotic organisms, rendering presence of this kinase exclusive to eukaryotes^26^.

**Figure 1.**
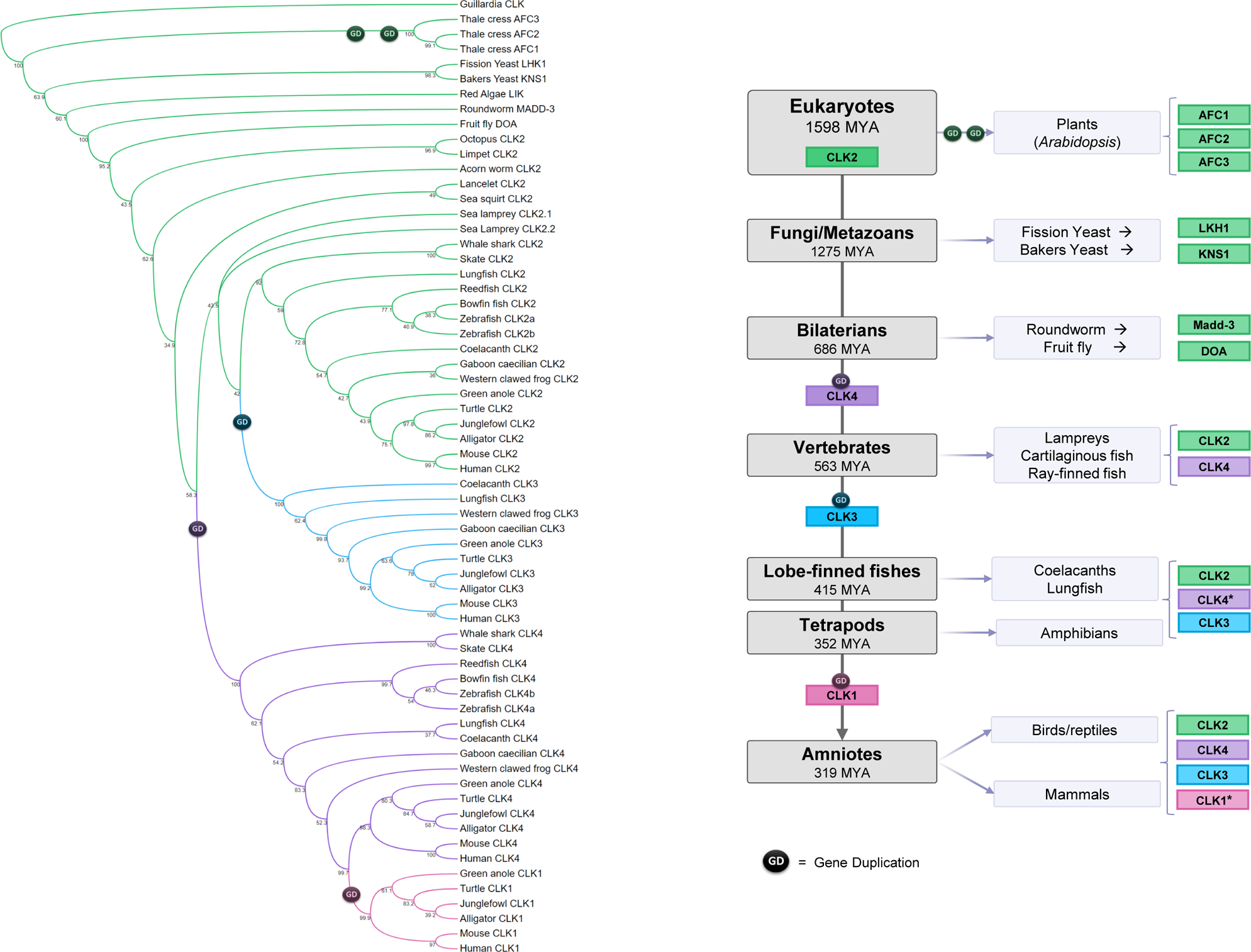
Phylogenetic analysis of the CLK family across eukaryotes. Phylogenetic tree of the CLK family (left) and a depiction of CLK gene expansion (right) in eukaryotic evolution. Homologous CLK proteins from eukaryotic species were identified using NCBI blastp. Kinase domain sequences were aligned using MAFFT, and a Neighbor-Joining phylogenetic tree was constructed with 1000 bootstrap replications. Branches are colour-coded according to their human paralog: CLK2 (green), CLK4 (purple), CLK3 (blue), CLK1 (pink). (* Two instances of gene loss: CLK4 in Neobatrachia and CLK1 in Psitacopasserae lineages. Refer to Fig. S3 for details)

Several duplication events leading to the production of paralogs were identified (Figure 1). Firstly, within *Arabidopsis thaliana* (with plants being the most distant eukaryotic lineage to humans), two duplications of the original CLK2-related gene occurred, resulting in the presence of three CLK paralogs which are unique to plants (AFC1-3, Figure 1). Following the divergence of plants, three critical duplication events appear to account for the four CLKs found in mammals. This includes **(1)** the original CLK2 orthologous gene being duplicated in the ancestor of all vertebrates to create the first CLK4 ortholog, **(2)** another duplication of the CLK2 ortholog, giving rise to CLK3-orthologous genes in the lobe-finned fishes, and finally, **(3)** duplication of the CLK4 ortholog to create the first CLK1 gene, present only in amniotes (Figure 1). In addition to this, within the teleost lineage of ray-finned fishes (e.g. Zebrafish), an additional gene duplication of CLK2 and CLK4 was observed. This is likely attributed to the well-known whole genome duplication event that occurred in the common ancestor of all extant teleosts^27,28^. Indeed, the presence of only one copy of CLK2 and CLK4 in Bowfin (*Amia calva)* and Reedfish (*Erpetoichthys calabaricus*); (fish species that diverged just prior to teleosts), support this conclusion.

Interestingly, each of the three CLK duplication events identified in vertebrates align with significant milestones in the evolution of eukaryotic complexity. The initial duplication produced CLK4 in the sea lamprey *Petromyzon marinus*, which coincides with a crucial whole genome duplication event at the baseline of all vertebrates^29^. The second duplication produced CLK3 in lobe-finned fishes, an intermediate lineage between ray-finned fishes and amphibians, a lineage representing the transition from water-to-land^30,31^. Following this, complete eukaryotic terrestrialisation was completed during the evolution of amniotes, corresponding to the third duplication which introduced CLK1. Additionally, CLK gene expansion occurred in plants sometime during their terrestrialisation. Phylogenetic analysis revealed only one CLK2-related gene in the water-living Chlorophyta, *Chlamydomonas reinhardtii*, while additional homologs are present in all land-dwelling plants, a transition that would have required extensive adaptation^10^.

Previous studies have demonstrated that as organisms evolve towards greater complexity, there is a corresponding rise in both the intricacy of AS and duplications of splicing regulator genes^32–34^. This is because the increased regulation of AS, such as that by the CLK family, offers a source of transcriptional diversity to facilitate adaptation. While there is general conservation of core spliceosomal proteins, there is a selective expansion of protein families in metazoans that are involved in splicing regulation, including vertebrate-specific duplications of hnRNPs and SRPKs^34^. As such, these CLK duplication events likely reflect the increase in splicing complexity necessary for the corresponding shifts in eukaryotic evolution.

As a final comment, it is important to note that we have identified various database gene annotation errors where CLK4 is incorrectly labelled as CLK1, likely due to their high similarity. These errors are easily identifiable by the presence of “CLK1” within species that diverged prior to amniotes, the lineage where CLK1 originated. We encountered only one publication that appears to have used one of these incorrectly named CLKs. In this study, they characterised an embryonic knockdown of “CLK1” in *Xenopus tropicalis*, demonstrating minor phenotypic changes, including a bent axis at the late tailbud stage^8^. However, our phylogenetic analysis reveals that this gene is orthologous to CLK4, not CLK1, which could impact how these results are interpreted in the context of mouse models.

#### Gene preservation and gene loss

It is widely acknowledged that genes deemed functionally indispensable are less likely to be lost during evolution and therefore retained within the genome^35^. Despite the prevalence of CLKs throughout eukaryotes, some single-cell organisms have lost CLK genes entirely^25^. Moreover, studies involving genetic knockout of the sole CLK homolog in six different yeast species demonstrate viability, further demonstrating that some single-celled organisms can survive without this kinase family^11,36–41^. On the contrary, there are no documented instances of natural CLK gene loss within multicellular eukaryotes. Additionally, genetic knockout of the well-studied CLK2 ortholog “DOA” (*Drosophila melanogaster,* fruit fly) results in severely abnormal neural development and embryonic lethality^4,42^. This underscores the potential indispensability of these kinases in more complex organisms. Given these findings, we investigated whether natural CLK gene loss has occurred within metazoans to ascertain whether this gene family may be essential for complex multicellular life.

For this analysis, we used protein sequences of the kinase domains from human CLK1, CLK2, CLK3 and CLK4 to conduct blastp search across all (well annotated) metazoans to ascertain presence or absence (Supplementary Table 2). Our analysis, which included both the “non-vertebrate” and “vertebrate” categories, found no evidence of complete absence. Notably, all metazoan species belonging to the non-vertebrate category (which is the vast majority) were found to possess a gene orthologous to CLK2. While not definitive, the persistence of this gene implies essential functions for the CLK family within the context of metazoan life.

To further this analysis, we looked deeper into the vertebrate lineage to identify loss of individual CLK paralogs. To uncover genuine gene loss events and exclude the potential for incomplete/incorrect metagenome assemblies, we identified whether ancestral CLK paralog loss had occurred, i.e. in multiple related species, and excluded species-specific loss^43^. Through this analysis, we identified two instances of CLK gene loss within diverse, species-rich vertebrate lineages. The first pertains to the loss of CLK4 in Neobatrachia, a suborder encompassing over 96% of extant species of amphibians^44^, while the second involves the loss of CLK1 in the avian taxon Psittacopasserae, comprising over 60% of all bird species (Supplementary Figure 3, Supplementary Table 2). The absence of these genes suggests a lack of biological necessity for CLK1 or CLK4 within these vertebrate branches, which may have occurred in different ways. For example, during evolution, genes can lose functionality, rendering them dispensable which leads to their natural loss within the genome. Conversely, gene loss can occur abruptly and provide an adaptive advantage, securing this genetic variation in populations^35^.

Importantly, we found no instances of CLK2 or CLK3 gene loss, implying that they may be functionally indispensable CLK paralogs. Presently, complete transgenic knockout mice have only been generated for CLK1 and CLK2, both of which are shown to be viable and fertile in a controlled laboratory environment^45–48^, indicating that despite the preservation of CLK2 across metazoans, mice can survive and reproduce without this gene. One possible explanation is that while essential genes typically support growth and reproduction, “gene essentiality” is context dependent. Some genes are directly essential, impacting an organism’s fertility or viability upon removal, while others are indirectly essential, affecting long-term survival^49–51^. As such, CLK2 knockout may not immediately compromise survival in mice, but may decrease long-term survival fitness in natural environments. Alternatively, multiple CLK paralogs within an organism could have redundant functions and compensate in the absence of one another.

Although CLK3 has not been knocked out in a mouse model, a compelling study has explored the effect of embryonic knockdown of CLKs in the frog *Xenopus tropicalis*. This species possesses CLK2, CLK3 and CLK4 (mislabelled as “CLK1”) which are co-expressed in neural tissue during early embryogenesis^7,8^. Individual knockdown using translation-blocking morpholino oligonucleotides demonstrate CLK3 is the only ortholog essential for development, leading to a significant reduction in head and eye size. Embryos with a greater CLK3 knockdown efficiency experienced lethality, while knockdown of CLK2 and CLK4 produced only mild phenotypic changes in embryonic development^8^. This study is the first to establish an individual CLK paralog as essential for vertebrate life. Furthermore, this severe neural development phenotype shares common features with the embryonically lethal knockdown of DOA in *D. melanogaster*, which could implicate a conserved role^4,42^. Despite DOA being an ortholog of CLK2, a plausible explanation could be that when CLK3 originated from the duplication of CLK2, it acquired this developmentally essential function through the process of gene subfunctionalisation.

### (2) Structural comparisons

#### The conserved kinase domain

To gain insight into how this kinase family has adapted and diversified, and whether functions have remained conserved, we have compared the structural features of CLK paralogs (human CLK1-4, unique genes derived from duplications), and CLK orthologs (CLK2-related genes, diversified through speciation). All CLK homologs are composed of two primary regions: the kinase domain and the intrinsically disordered N-terminal. While the disordered nature of the N-terminal regions prevents structural analysis, we present, for the first time, a predicted structural alignment of the kinase domains of CLKs from various eukaryotic organisms. In addition, while crystal structure comparisons of human CLK1-4 kinase domains have been shown elsewhere^1,2,16,17^, we expand on these by connecting their sequence alignment to their specific structural features (Figure 2A-C).

**Figure 2.**
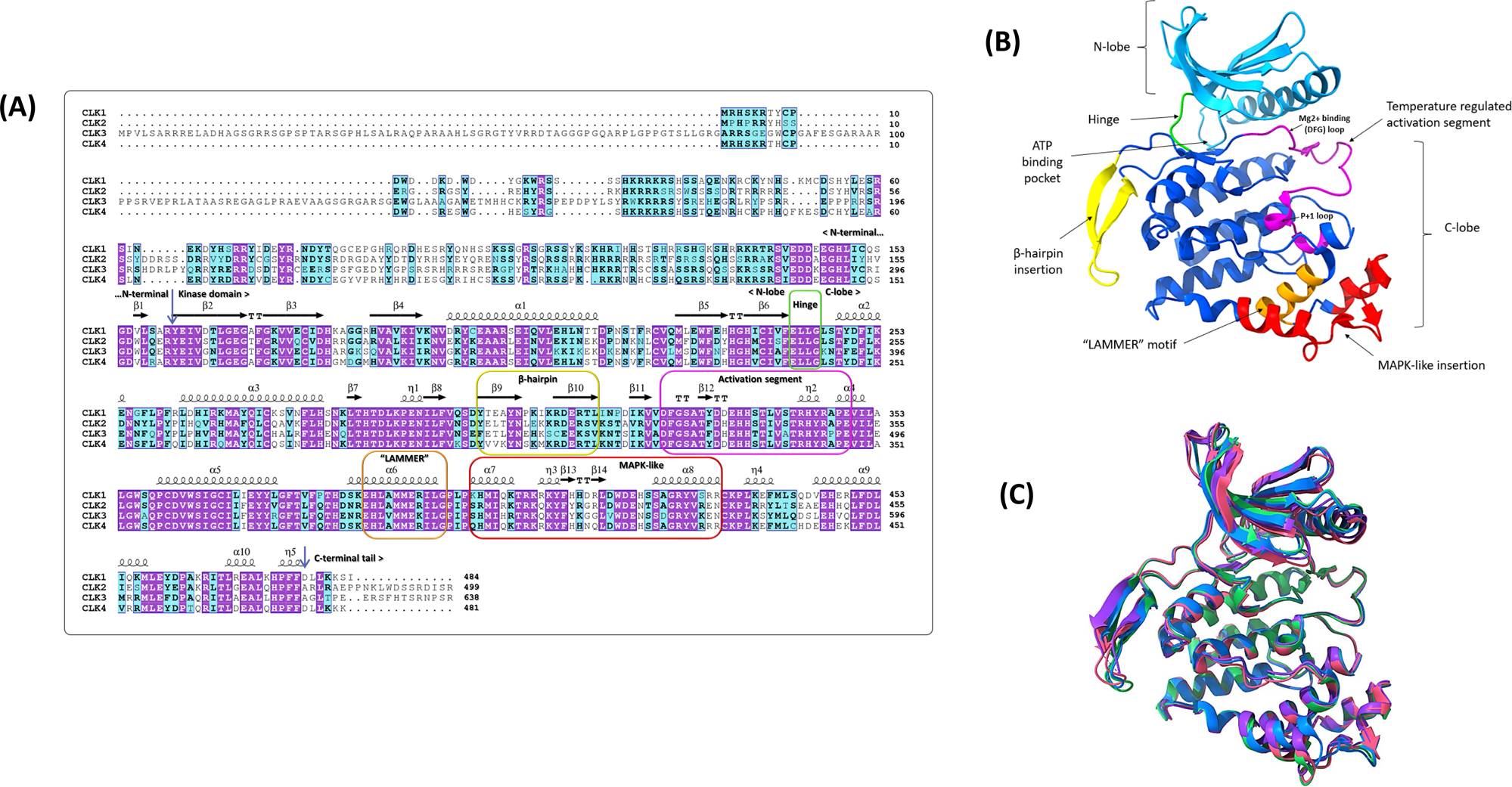
Sequence and structural alignment of human CLK1-4. **(A)** Amino acid sequences for human CLK1-4 were aligned using muscle, with conserved structural elements depicted above the alignment. Purple boxes denote strict identity, while blue boxes indicate 75% group similarity, with bold characters representing amino acids with similar physicochemical properties. α-helices and 3_10_-helices (η) are displayed as big and small squiggles, respectively. β-strands are shown as arrows, and strict β-turns as “TT” leters. **(B)** Crystal structure of CLK2 (PDB:6FYL) kinase domain with structural elements coloured and labelled corresponding to those in (A). **(C)** Crystal structures of CLK1-4 kinase domains (CLK1 PDB:6R8J - pink, CLK2 PDB:6FYL - green, CLK3 PDB:6Z53 blue, CLK4 PDB:6fyv - purple).

Human CLK1-4 demonstrate a high degree of conservation of the kinase domain and divergence of their N-termini, as shown by a sequence alignment of human CLK1-4 (Figure 2A), along with the kinase domain structures of CLK2 (Figure 2B) and CLK1-4 overlayed (Figure 2C). The enzymes display typical kinase features, including an ATP binding pocket (Figure 2B) situated within a hinge region (Figure 2A/B, green) linking the N- and C-lobes of the protein. The N-lobe comprises six β-strands and one α-helix, while the C-lobe is made up of fourteen helices (α or 3_10_), a β-hairpin, and six short β-strands (Figure 2A/B). Within the α6 helix, which is located within the C-lobe of CLK1-4, is the highly conserved “EH<u>LAMMER</u>ILG” motif (Figure 2A/B, orange). Preceding this motif is a distinct MAPK-like insertion which keeps the α6 helix inaccessible to solvents (Figure 2A/B, red)^52^. Additionally, this group of enzymes possess a unique insertion at the beginning of the C-lobe, forming an extended β-hairpin structure (Figure 2A/B, yellow). The C-lobe of CLKs contain the activation segment (Figure 2A/B, represented in magenta) positioned in front of the ATP binding pocket, which, as discussed below, is subject to unique temperature regulation.

We have compared the amino acid sequences (Table 1, Supplementary Figure 1) and AlphaFold structures (Figure 3) of nine CLK2 orthologs from diverse eukaryotes. Overall, the amino acid alignment indicates moderate conservation of the kinase domain across species, while the N-terminal region exhibits high variability (Supplementary Figure 1). The similarity of the kinase domain compared to human CLK2 decreases with greater evolutionary distance, yet it still retains significant conservation, with the ortholog AFC2 in plants sharing 48% identical amino acids (Table 1). Certain regions, including the “LAMMER” motif and activation segment, are highly conserved among species; however, the MAPK-like insert and β-hairpin show low sequence conservation (Supplementary Figure 1). Despite these differences, a high level of structural conservation is observed in the kinase domains of eight out of the nine CLK2 orthologs (Figure 3A) including human **CLK2**, *Mus musculus* (mouse) **CLK2**, *Xenopus tropicalis* (western clawed frog) **CLK2**, *Danio rerio* (zebrafish) **CLK2b**, *Arabidopsis thaliana* (plant) **AFC2**, *Caenorhabditis elegans* (roundworm) **MADD-3**, *Drosophila melanogaster* (fruit fly) **DOA**, and *Schizosaccharomyces pombe* (fission yeast) **LKH1**. This structural similarity is consistent with the conserved role of CLKs in regulating alternative splicing (AS) through the phosphorylation of SR proteins, a function confirmed in mammals, *D. melanogaster*^18,53,54^, and the most distant eukaryotic lineage of plants in *A. thaliana*^10,55^.

**Figure 3.**
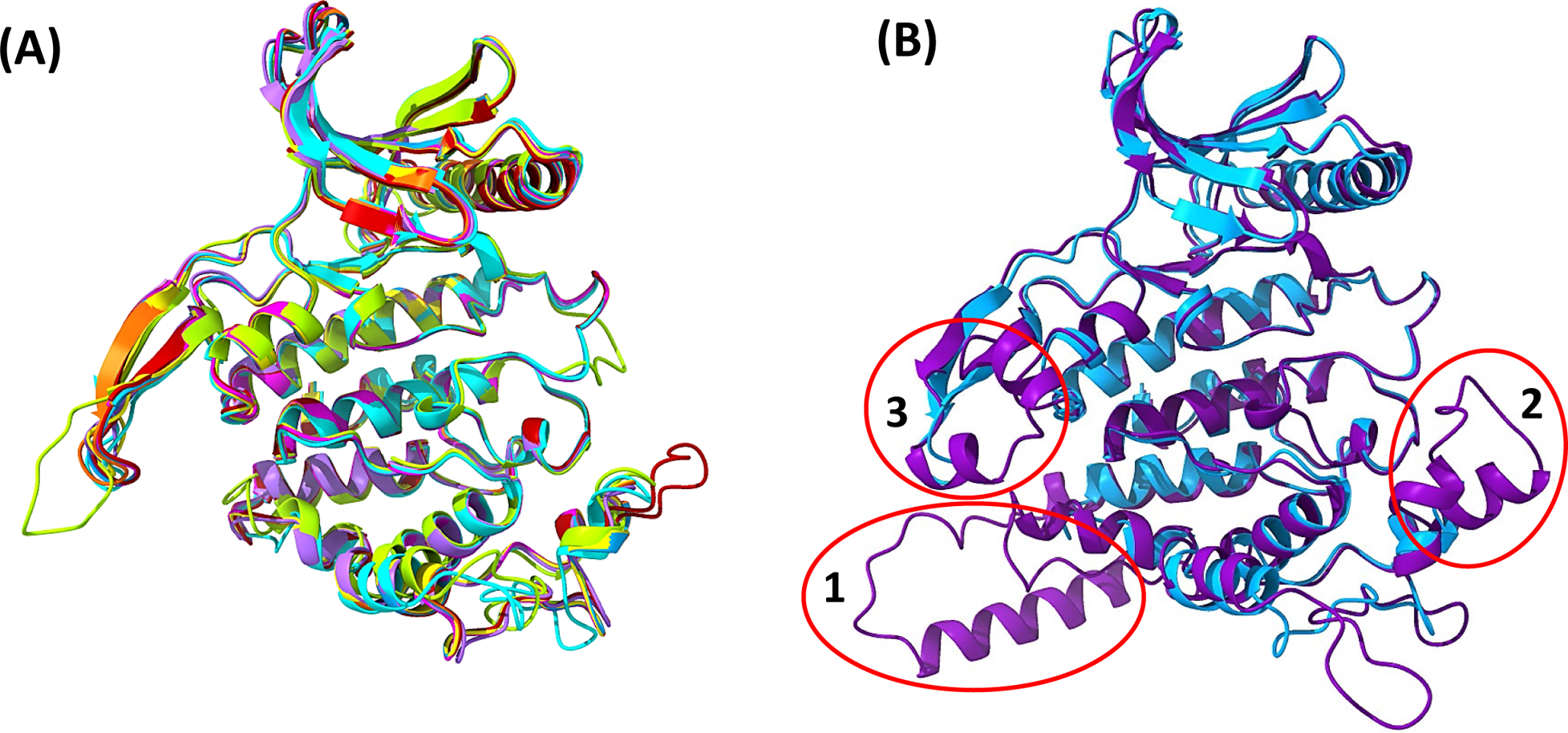
AlphaFold structural predictions of CLK2 orthologs. **(A)** AlphaFold structural overlays of the kinase domains of eight diverse eukaryotes: Human “CLK2” (blue), Mouse *“*CLK2” (dark orange), Western clawed frog - *Xenopus tropicalis “*CLK2” (yellow), Zebrafish - *Danio rerio “*CLK2b” (fuchsia), Plant - *Arabidopsis thaliana “*AFC2” (lime green), Roundworm - *Caenorhabditis elegans “*MADD-3” (light purple), Fruit Fly - *Drosophila melanogaster “*DOA” (red) and Fission yeast - *Schizosaccharomyces pombe “*LKH1” (cyan). AlphaFold structures were downloaded from UniProt, and terminal ends were removed to overlay the kinase domains. **(B)** AlphaFold overlay of Bakers yeast - *Saccharomyces cerevisiae “*KSN1” (dark purple) with Human “CLK2” (blue) showing unique inserted structural elements, three major additions are circled in red.

**Table 1:** Percent similarity of aminos acid sequences from CLK2 orthologs across eukaryotes.

| Human CLK2 orthologs |  | Whole protein | Kinase domain |
| --- | --- | --- | --- |
| Mouse | CLK2 | 50% | 98% |
| Western clawed frog | CLK2 | 50% | 90% |
| Zebrafish | CLK2b | 48% | 84% |
| Fruit fly | DOA | 29% | 72% |
| Roundworm | MADD-3 | 24% | 63% |
| Plant | AFC2 | 33% | 48% |
| Fission Yeast | LKH1 | 24% | 44% |
| Baker's Yeast | KNS1 | 23% | 31% |

Our comparisons have revealed an outlier, indicating a significant structural variation that has arisen during yeast diversification. Despite the conservation of LKH1 from the fission yeast *S. pombe* (Figure 3A, cyan), a second yeast species, *S. cerevisiae*, has three unique regions within its CLK2 ortholog **KNS1** (Figure 3B, purple). These include (i) a large helical insertion following the MAPK-like section (Figure 3B, red circle #1), (ii) a smaller insertion within the MAPK-like section (Figure 3B, red circle #2), and (iii) a helical section at the tip of the β-hairpin (Figure 3B, red circle #3). Interestingly, the differences between KNS1 and LKH1 may reflect the distinct splicing mechanisms present in *S.* c*erevisiae* and *S. pombe*, as well as the functional roles of these kinases. While research suggests that spliceosomal complexity was high in the LECA, it underwent simplification in many eukaryotic lineages, including during the evolution of yeast^32,56,57^. In the case of *S. cerevisiae*, this yeast demonstrates low usage of splicing, with only 4% of protein coding regions requiring constitutive splicing and an even lower percentage (0.2%) undergoing AS; typically through intron retention^58,59^. Furthermore, knockout of KNS1 from *S. cerevisiae* does not change the ratio of pre-mRNA to mRNA^60^, demonstrating this CLK2 ortholog is not involved in either constitutive splicing or AS. In stark contrast, *S. pombe* demonstrate greater diversity of splicing, with 43% of genes containing one or more introns, and 4.5% of genes undergoing AS^61^. Furthermore, unlike KNS1, knockout of LKH1 (Figure 3A, cyan) increases the ratio of pre-mRNA to mRNA by 8-fold^62^. However, this genetic alteration does not alter global AS patterns^63^. This suggests that LKH1 is essential for constitutive splicing but does not play a significant role in AS, exemplifying another instance of yeast simplification of splicing mechanisms. Therefore, in the absence of AS regulation, KNS1 has adapted these unique structural elements to potentially govern different biological functions. For example, KNS1 has functional roles in regulating various stress responses in *S. cerevisiae*. These include high pH stress^60^, nutrient limitation, and rapamycin treatment^64^.

It is intriguing that despite not being involved in AS regulation, LKH1 exhibits high structural conservation with other CLK2 orthologs (see Figure 3A). Since these kinases are primarily known to regulate splicing through the phosphorylation of SR proteins, it is tempting to speculate that this shared structure is crucial for targeting these proteins. This interaction is known to occur in *S. pombe*, where the structurally conserved LKH1 phosphorylates its SR protein, Srp1, *in vivo*^65^. In contrast, yeast two-hybrid screens have indicated that KNS1, which lacks structural conservation, is still capable of interacting with and phosphorylating SR proteins from other species^66^. As such, it appears the non-conserved structural elements in KNS1 do not impair its ability to phosphorylate SR proteins *in vitro*.

These findings highlight the structural and functional diversification of CLKs in yeast species, offering potential benefits for research applicable to practical fields. For example, CLK homologs are known to regulate the pathogenicity of disease causing yeast species in humans^39,40^ and crop^41^ species. Additionally, these kinases are involved in yeast flocculation regulation, a key molecular mechanism for industrial fermentation^67,68^.

#### The diverged N-terminal

The N-terminal region of CLKs serves as a crucial regulatory mechanism for kinase function and specificity towards SR proteins. An essential property of this domain is its intrinsically disordered and unstructured nature. To assess whether this feature is conserved, we assessed protein disorder of CLK homologs across diverse eukaryotes. We found that, consistently, these kinases are divided by a highly disordered N-terminal followed by an ordered kinase domain (Supplementary Figure 2). Disordered regions are typically diverged in their amino acid composition across species, yet they maintain other conserved properties that enable them to perform similar biological functions^69^. Accordingly, CLKs across eukaryotes have retained the ability to phosphorylate SR proteins^18,19,65^, despite highly variable N-termini (see Supplementary Figure 1). In addition, expression of the N-terminal alone can bind to SR proteins, even those originating from diverse organisms, highlighting a conserved function^66^. However, divergence in N-termini among CLK homologs facilitates distinct regulation of substrate specificity, kinase activity, and subcellular localisation

An important variable between the N-termini of CLK homologs is the adaptation of Arginine-Serine (RS) motifs, which are more prevalent in higher order metazoans. The expansion of these motifs forms an “RS domain”; a critical feature which is also present in their SR protein substrates. In organisms that possess a higher abundance of RS motifs, such as mammals, this allows for self-association, leading to oligomerization^70^. As a consequence, oligomerized CLK1 has a greater affinity for its SR protein substrate SRSF1, which in turn, enables phosphorylation of 18 serine residues on the SR protein. In contrast, expression of N-terminally truncated CLK1 only allows for phosphorylation of 6 serine residues within SRSF1, demonstrating this region is critical for RS domain hyperphosphorylation^71^.

CLK homologs are also regulated through autophosphorylation of their N-termini to mediate kinase activity. This regulatory function is maintained in diverse organisms, including yeast, mammals, plants and fruit fly, however, the phosphorylated sites are highly variable between CLKs^14,19,64,72–74^. It also appears that not all CLKs undergo autophosphorylation, such as AFC1 in *A. thaliana*. In contrast, the paralog AFC2 within this organism is autophosphorylated, which activates its ability to interact with the SR protein, SR33, demonstrating distinct regulatory mechanisms between these CLKs^19^. Furthermore, individual CLKs undergo differential phosphorylation to control their activity and specificity. Research by Prasad and Manley ^75^ found that the pattern of autophosphorylation on CLK1 regulates kinase activity and specificity towards its SR protein substrates. Herein, they demonstrated that (1) autophosphorylation of CLK1 on tyrosine residues (but not serine/threonine) dictates specificity towards SRSF1, (2) autophosphorylation of CLK1 on serine/threonine residues dictates specificity towards SRSF2, and (3) phosphorylation of SRSF5 remains unaffected by the pattern of CLK1 autophosphorylation. In addition to this, autophosphorylation of CLK2 on Ser141 has been shown to affect its subnuclear localisation, promoting translocation from nuclear speckles to the nucleoplasm^76^.

The importance of the N-terminal regarding subcellular localisation has been highlighted in two model organisms, namely *D. melanogaster* and *C. elegans*. The CLK2 orthologs within these organisms are alternatively spliced to produce either nuclear-specific, or cytoplasmic-specific isoforms, with the only distinguishing factor being a difference in their N-termini^77–79^. In both species, these distinct isoforms perform unique functions within their specific subcellular compartments. Coupled with this, in mammals, truncation of CLK1’s N-terminal causes cytoplasmic accumulation of the kinase^80^. What is particularly intriguing about this observation is that CLK1 lacks a functional nuclear localisation signal. Consequently, the protein depends on the interaction between its N-terminal and its substrate, SRSF1, forming a stable complex. Following phosphorylation, this complex binds to SRSF1’s nuclear importer, TNPO3, facilitating CLK1 translocation from the cytoplasm^80^.

### (3) Unique thermo-regulation of CLKs

#### The temperature-regulated activation segment

A remarkable property of the CLK family is that their activity is upregulated by slight decreases in the physiological growth temperature of an organism. So far, this temperature regulation has been observed in CLK homologs from diverse eukaryotes, including those found in mammals^6^, *Drosophila melanogaster*^6^, *Arabidopsis thaliana*^10^, *Trachemys scripta*^6^, *Alligator mississippiensis*^6^ and *Cyanidioschyzon merolae*^23^ (Figure 4B, marked with Asterixis). Groundbreaking research by Haltenhof, et al. ^6^ discovered that this regulation occurs within the activation segment of the kinase domain. However, no report has investigated how the temperature-regulated activation segment has functionally adapted and diversified throughout eukaryotic evolution.

**Figure 4.**
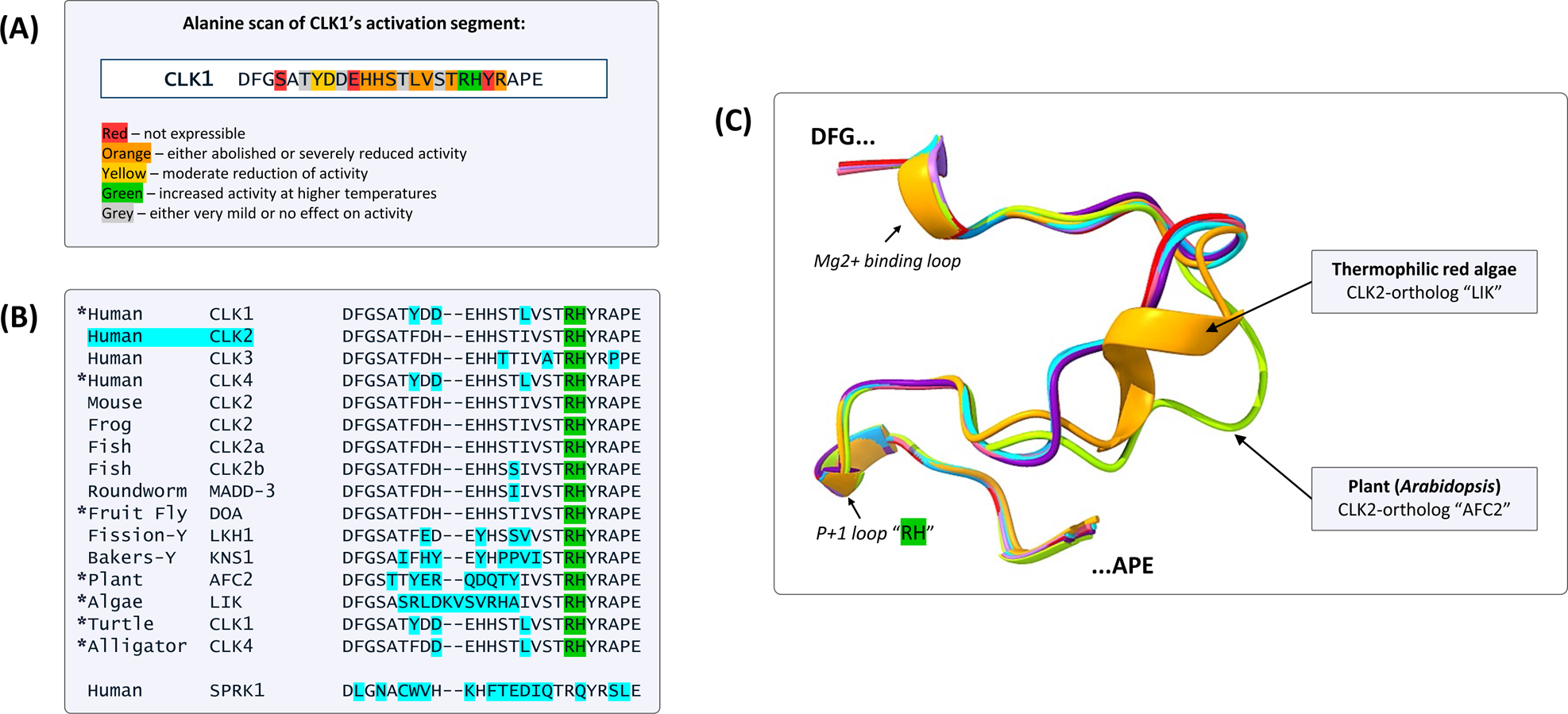
Alanine scan and sequence/structural alignment of the temperature regulated activation segment in CLK homologs. **(A)** Summary of the alanine scan performed by Haltenhof et al. (2020). Individual amino acids within the activation segment of CLK1 were substituted for alanine to assess the impact on kinase activity across a temperature gradient from 33°C to 40°C. Results were interpreted and highlighted accordingly. Additionally, the substitution of the green “H” (Histidine H343) with a “Q” (Glutamine) abolished temperature sensitivity. **(B)** Amino acid alignment of kinase activation segments of CLK homologs and human SRPK1. Sites highlighted blue represent positions that differ from human CLK2. Asterixis denote kinases confirmed to be thermosensitive. **(C)**AlphaFold structures of CLK homolog activation segments from various organisms: Human/mouse - “CLK1/CLK4” (pink) and “CLK2” (blue), Roundworm - *Caenorhabditis elegans “*MADD-3” (light purple), Fruit Fly - *Drosophila melanogaster “*DOA” (red), Plant - *Arabidopsis thaliana “*AFC2” (lime green), Fission yeast - *Schizosaccharomyces pombe “*LKH1” (cyan), Bakers yeast - *Saccharomyces cerevisiae “*KSN1” (dark purple) and Thermophilic red algae - *Cyanidioschyzon merolae* “LIK”.

Like other kinases, the activation segment of CLK homologs is located in front of the ATP binding pocket (Figure 2B, magenta). Herein, the ATP-magnesium counter-ion directly interacts with the DFG loop, facilitating the transfer of the γ-phosphate to specific substrates. Within the activation segment is the P+1 loop, which serves as an important site for substrate interaction (Figure 2B, 4C). This domain undergoes a conformational change to regulate accessibility and correct positioning of the nucleotide which, for many kinases, is brought about through phosphorylation of this region^81^. However, in the case of CLKs, the activation segment is thermally-regulated, typically in a reversible manner^6^. This regulation is particularly unique for CLKs, which follow an opposite trend to a standard enzymatic Q_10_ temperature coefficient, which is a calculation of the thermo-sensitivity of the reaction rate. For most mesophilic enzymes, Q_10_ = 2-3, meaning that for each 10°C rise above their physiological temperature, their reaction rate doubles or triples^82^. Comparatively, CLK1 and CLK4 in humans have a negative Q_10_ value, demonstrating minimal kinase activity at 38°C and ∼4-fold increase in kinase activity when reduced to 35°C.

The activation segment is a crucial element for thermo-regulation of CLKs. Exchanging this domain in CLK1, with that of the temperature insensitive SR protein kinase 1 (SRPK1), leads to loss of CLK1’s temperature regulation and a gain of activity at higher temperatures *in vitro*^6^. Likewise, the addition of CLK1’s activation segment to SRPK1 leads to induced temperature-sensitivity of the latter. To identify the specific thermo-sensitive region within the activation segment, an Alanine scan has been performed (Haltenhof, et al. ^6^) and kinase activity measured within the temperature range of 33-40°C (summarised in Figure 4A). The replacement of arginine (R342) and histidine (H343), two highly conserved adjacent amino acid residues located within the P+1 loop (highlighted in green, Figure 4A-C), led to changes in the ability of temperature to regulate activity. In this case, an increase in kinase activity was shown at higher temperatures. Furthermore, exchanging H343 with the corresponding amino acid in SRPK1, glutamine (Q516), led to a gain of temperature sensitivity in SRPK1 and complete loss of thermo-regulation of CLK1^6^. Thus, it is evident that H343 plays a significant role in conferring the ability of the kinase to respond to temperature, with a single amino acid substitution having the potential to transform a non-thermally regulated kinase into a thermally regulated one.

#### Species-specific temperature regulation

The sequence (Figure 4B) and structural (Figure 4C) alignment of CLK activation segments from diverse eukaryotes reveals a mix of both conserved and non-conserved features. Compared to human CLK2, amino acid alignment shows the middle region of the activation segment to be less conserved in more diverged species, while the terminal regions exhibit high conservation (blue, Figure 4B). Despite these differences, temperature regulation has been confirmed in CLK homologs across diverse species (Asterixis, Figure 4B). In accord with this, there is strict conservation of the thermo-sensitive “RH” motif within the P+1 loop (green, Figure 4B). As such, it is predicted that temperature regulation of CLKs is a function of ancient origin, likely occurring in the LECA. Structural comparisons further demonstrate that this segment is mostly conserved. In agreement with their known temperature regulation, mammalian CLK1/CLK4 and *D. melanogaster* DOA share structural conservation of their activation segments (Figure 4C). Additionally, mammalian CLK2, *C. elegans* MADD-3, *S. pombe* LKH1, and even the diverged KNS1 from *S. cerevisiae*, share this structural conservation, indicating that these kinases, although not formally shown, are also likely temperature dependent (Figure 4C).

In contrast, two CLKs demonstrate structural divergence of their activation segments despite their known thermo-sensitivity properties, including *A. thaliana*, AFC2 (Figure 4C, lime green), and *C. merolae*, LIK (orange, Figure 4C)^6,23^. In both cases, this could explain their unique temperature regulation profiles. Firstly, AFC2 is the only reported CLK homolog that is irreversibly inactivated by temperature^10^. The activation segment of AFC2 contains 9 non-conserved amino acids (Figure 4B), with structural predictions demonstrating a change in orientation (Figure 4C, lime green). However, to date, it is unknown how this irreversible inactivation is achieved through the activation segment. Secondly, *C. merolae* is a unicellular red algae that grows in hot springs up to 56°C. Its CLK2 ortholog, LIK, has adapted as a thermophilic enzyme and has been suggested to reach maximum activity at 48°C, and inhibition at 56°C. Sequence alignments show LIK contains 11 non-conserved amino acids (Figure 4B), including a two amino acid insertion within its activation segment. Structural comparisons suggest that these alterations result in the formation of an α-helix (Figure 4C, orange), which as a secondary structure, may possess stability at higher temperature. This property potentially enables LIK to be regulated at higher temperatures, assisting *C. merolae* adaption in extreme conditions.

The adaptation of an organism-dependant temperature profile is not limited to LIK, but is a facet seen for all other CLK homologs so far tested^6,10,23^. For example, in reptiles such as turtles and alligators, CLK1/CLK4 show temperature regulation with maximal activity at 25°C and inhibition at 35°C. Meanwhile, DOA (*D. melanogaster*) is inhibited at 32°C, and maximal activity is seen at 20°C. Human CLK1 and CLK4, in contrast, experience a significant activity change within their physiological temperature range of 33-38°C, with maximal activity between 22-24°C^6^. Interestingly, unlike the case of LIK, the variation in temperature profiles cannot be explained by structural (Figure 4C – human vs *D. melanogaster*) or sequence (Figure 4B – turtle vs human CLK1) differences in the activation segments. This indicates other regions within the kinase contribute to species-specific temperature profiles. A possible contributing factor is the disordered N-terminal^6^. Deletion of N-termini of both human CLK1 and CLK4 leads to a reduced temperature-activity profile compared to the full-length proteins. Therefore, while the activation segment may not entirely dictate the precise temperature range of activity, it appears to be a critical contributor to this process.

#### Temperature controlled biological functions

The unique thermosensitivity of CLKs adds an intriguing dimension to their role in the regulation of alternative splicing (AS). Aside from generating novel protein isoforms, AS can modulate gene expression by producing “poison” transcripts that undergo mRNA decay^83^. It is through this mechanism that body temperature controlled AS profoundly shapes global gene expression, having evolutionary roots deeper than the core circadian clock itself. Of significance, the family of SR proteins across diverse eukaryotes are subject to regulation in this manner, modulating their expression in a temperature-dependent fashion and establishing a feedback loop for AS-linked mRNA decay^83^. CLKs influence a large proportion of the temperature-dependant transcriptome, regulating over 50% of thermo-sensitive exons *in vitro*^6,10,24^. Physiological functions linked to the temperature control of CLKs include circadian rhythm oscillations^6,24^, reptilian temperature-dependent sex determination (TSD)^6^ and plant thermomorphogenesis^10^.

Circadian rhythms operate on a ∼24hr cycle, coordinating the cyclic expression of genes to regulate organism physiology over the course of a solar day. These systems are observed across diverse organisms, spanning from mammals to plants, with light and temperature serving as two major universal timing cues^84^. In response to light, mammals adjust their body temperature by 1-4°C in day-night cycles, which is accompanied by rhythmic AS through altered phosphorylation of SR proteins^24,85–87^. For instance, the expression of Cold-inducible RNA-Binding Protein (CIRBP) oscillates in a temperature-dependent manner, regulating many important circadian mRNAs, including “CLOCK”^87^. This rhythmic expression of CIRBP is controlled by CLK-dependent AS via differential exon inclusion generating a poison transcript^6,88^. In mice, decreased core body temperature (day) results in exon 7a inclusion, producing the full-length CIRBP transcript, whereas increased core body temperature (night) leads to exon 7b/8 inclusion, triggering nonsense-mediated decay.

Due to their conserved thermosensitivity, it is interesting to consider that CLKs in diverse eukaryotes may also play a role in regulating the circadian rhythm. In addition to mammals, temperature-dependant AS of circadian genes have been observed in fish^89^, yeast^90,91^, plants^92^, and fruit fly^93,94^. In *Drosophila*, the circadian clock is influenced by temperature-dependent AS of “TIM”^94^, a protein sharing homology with an isoform of mammalian U2af26 that includes exons 6/7 (U2af26Δ67)^95^. Both interact with and affect the stability of “PERIOD” homologs in their respective species, demonstrating shared functionality. Notably, U2af26Δ67 undergoes rhythmic splicing in a CLK-dependent manner due to circadian temperature fluctuations^86^. As such, it is tempting to speculate that the CLK2-ortholog, DOA, might participate in temperature-mediated AS of TIM to regulate the circadian cycle in *Drosophila*.

Across diverse reptiles, sex is determined by the temperature at which their eggs are incubated. In the turtle *Trachemys scripta elegans*, embryonic development at 26°C produces all males, whilst those incubated above 31°C, produces all females. At temperatures in between, the broods will give rise to individuals of both sexes. Although the mechanisms of TSD are not well understood, researchers have suggested that AS of the polycomb-repressive complex 2 component “JARID2” may play a role. Herein, males produced at 26°C preferentially retain intron 15. *In vitro* studies indicate that inhibiting CLKs at 26°C reduces intron 15 retention of JARID2, suggesting the involvement of this kinase family in TSD. Furthermore, CLK1 in *T. scripta* has full activity below 26*°*C, and significant (∼90%) reduction above 31°C, essentially representing an on-off switch for TSD in this species.

In *Arabidopsis*, the CLK2-ortholog AFC2 is involved in the regulation of thermomorphogenesis, a process through which plants respond to high temperatures by elongating their hypocotyls/petioles and elevating their leaves to assist in cooling^96^. Mutation of AFC2 or inhibition with TG003 negatively regulates this process, causing an exaggerated high-temperature phenotype and a significant reduction in temperature-dependant AS^10^.

In the fission yeast, *S. pombe*, deletion of the CLK2 ortholog LKH1 leads to temperature-dependant changes in poly(A)+ mRNA localisation^62^. In LKH1 knockout cells grown at 36°C, 23% of the cells exhibit nuclear accumulation of mRNA, while at 30°C, 70% of the cells show cytoplasmic clustering of mRNA. In contrast, the mRNA in wild-type cells grown at these temperatures is uniformly distributed throughout the nucleus and cytoplasm. This observation suggests CLKs regulate mRNA subcellular distribution, which could potentially occur through their SR protein substrates, which have a critical function in mRNA export^97^. In a second yeast species, *Candida albicans*, knockout of its CLK2 ortholog KNS1 revealed a role in dimorphic transitioning^39^. Interestingly, higher incubation temperatures, which would reduce KNS1 activity, are known to increase the occurrence of the yeast-to-hyphae transitioning, suggesting this kinase may regulate the morphogenesis of *Candida albicans* in a temperature dependent manner^98,99^. Finally, the Σ1278b filamentous strain of *S. cerevisiae*, exhibits temperature-sensitive defects in filamentous growth. Remarkably, this can be overcome by either knocking out KNS1, or by overexpressing individual genes downstream of the MAPK signalling pathway to activate filamentous growth^100^. As such, temperature dependent control of KNS1 activity may be a negative regulator of flocculation via the MAPK pathway in this species.

The physiological functions of CLK may be quite diverse, however the question of whether all the events are occurring through changes in AS or other mechanisms remains to be solved. What is clear is that CLKs regulate a significant portion of the temperature-dependent transcriptome. Therefore, while the full implications of the CLK family being thermo-sensitive are not yet fully understood, this area of research holds significant promise for new insights and innovation.

## Concluding remarks

This study expands on our understanding of the evolution and diversification of the CLK family. Their significance is emphasised by their prevalence throughout eukaryotes, with all metazoans possessing at least one member in this family. This suggests they are essential genes in more complex organisms. CLK homologs exhibit strong functional conservation in regulating alternative splicing, phosphorylation of SR proteins, and responding to temperature changes. However, they have also adapted functions such as N-terminal regulation and thermo-sensitivity in a species-specific manner to meet an organisms needs.

The intricate interplay between CLKs and their substrates orchestrates temperature-dependent gene expression programs, highlighting the complexity of eukaryotic regulatory networks. As we comprehend the molecular implications of the newly found temperature regulation of CLKs, we realise how much there is to uncover about their biology. Understanding how CLKs integrate environmental and internal cues to modulate gene expression programs will deepen our understanding of eukaryotic biology and evolution. The ability of the Last Eukaryotic Common Ancestor to respond to temperature likely contributed towards its ability to adapt and survive, shaping the dynamic landscape of gene expression in eukaryotes. As CLK research sheds light on thermo-regulatory mechanisms in organisms, this may offer insight into how they will respond to climate change, such as for reptilian TSD, which could aid environmental conservation efforts. Additionally, our findings have significant implications for understanding CLK biology in human health. The association of CLK dysregulation with various human diseases underscores the importance of further elucidating their functional roles and potential as therapeutic targets. Moreover, our study provides valuable insights for studying CLKs in model organisms and their relevance to human biology.

## Methods

### Phylogenetic analysis

For the construction of the phylogenetic tree, protein sequences of 64 CLK homologs from 28 different species (Supplementary Table 1) were utilised. Sequences of their kinase domain, with an additional 24 amino acids on the N-terminal end (to improve bootstrapping), were aligned using MAFFT^101^. A phylogenetic tree was generated using Geneious 2023.2.1 (https://www.geneious.com) employing the Neighbor-Joining method, the Jukes-Cantor substitution model, a 30% support threshold greedy clustering, and bootstrap testing with 1000 replications. The final tree was formatted using iTOL^102^.

### Gene preservation and gene loss

The amino acid sequences of human CLK kinase domains were NCBI blastp search against all metazoan species (that contain >1000 proteins) in the refseq protein database (retrieved September 2023). From here, the protein alignments showing less than 40% similarity were excluded and then isoforms originating from the same gene were grouped together based on their gene symbol for each refseq ID. The final output included a list of unique CLK genes for each metazoan species, along with their associated blastp matches (Supplementary Table 2). Every metazoan species searched possessed a minimum of one CLK homolog and there was no evidence of complete CLK gene family loss. For the investigation of paralog loss, specifically in vertebrates, the blastp output for each species was manually checked. This list of vertebrate species was uploaded to TimeTree^103^ and an evolutionary timescale phylogenetic tree was generated. The tree was formatted using iTOL^102^ to depict the presence and absence of CLK paralogs from our output.

### Structural comparisons and PondR Prediction

Crystal structures for human CLK1-4 were obtained from the Protein Data Bank (PDB), with the following accession codes: CLK1 (PDB: 6R8J), CLK2 (PDB: 6FYL), CLK3 (PDB: 6Z53), and CLK4 (PDB: 6FYV). AlphaFold predicted structures for other CLK homologs were acquired from UniProt (refer to Supplementary Table 1). Subsequently, the structures of the kinase domains or activation segments were extracted and superimposed using ChimeraX^104^. Additionally, sequences were run through PondR (Protein DisOrder prediction System)^105^ to determine disordered/ordered regions.

### Sequence alignments

Whole protein sequences were downloaded from either NCBI or Uniprot (Supplementary Table 1) and aligned using the MUSCLE algorithm^106^. For human CLK1-4, this alignment was uploaded to ESPript^107^ along with their PDB files to align structural features. All elements shown are conserved between CLK1-4 kinase domains, however, there were a couple of features did differ in length by just one amino acid.

## Notes

### Competing Interest Statement

The authors have declared no competing interest.

### Summary of Updates

Small changes to spelling/wording throughout text to improve clarity

