## Supplementary Figure legends for "Comparison of the CDC2-like kinase family across eukaryotes highlights the functional conservation of these unique biological thermometers"

**Figure S1. Alignment of CLK2 orthologs.**

CLK2 orthologs were aligned using the Muscle algorithm. Purple boxes denote strict identity, while blue boxes indicate 75% group similarity, with bold characters representing amino acids with similar physicochemical properties.

**Figure S2. CLK sequence disorder prediction using PondR.**

The full-length protein sequences of CLK homologs were analysed using PondR (Protein DisOrder prediction System), revealing disordered N-terminals and C-terminal tails, and ordered kinase domains across all organisms. The disorder probability graphs are depicted, with the x-axis representing amino acid number and the y-axis representing disorder probability. These probabilities were calculated with a 5% false positive rate, where values above 0.5 (red line) indicate disorder.

**Figure S3. Gene loss of the CLK family in vertebrates.**

CLK homologs were found across vertebrate species. A list of species were uploaded to TimeTree, and an evolutionary timescale phylogenetic tree was generated. In the tree, filled circles denote the presence of a CLK gene, empty circles signify the absence of a CLK gene (that would be expected based on phylogenetics), and stars indicate potential presence of additional paralogs.
