## Supplementary Figure 1 and 2 for "Comparison of the CDC2-like kinase family across eukaryotes highlights the functional conservation of these unique biological thermometers"

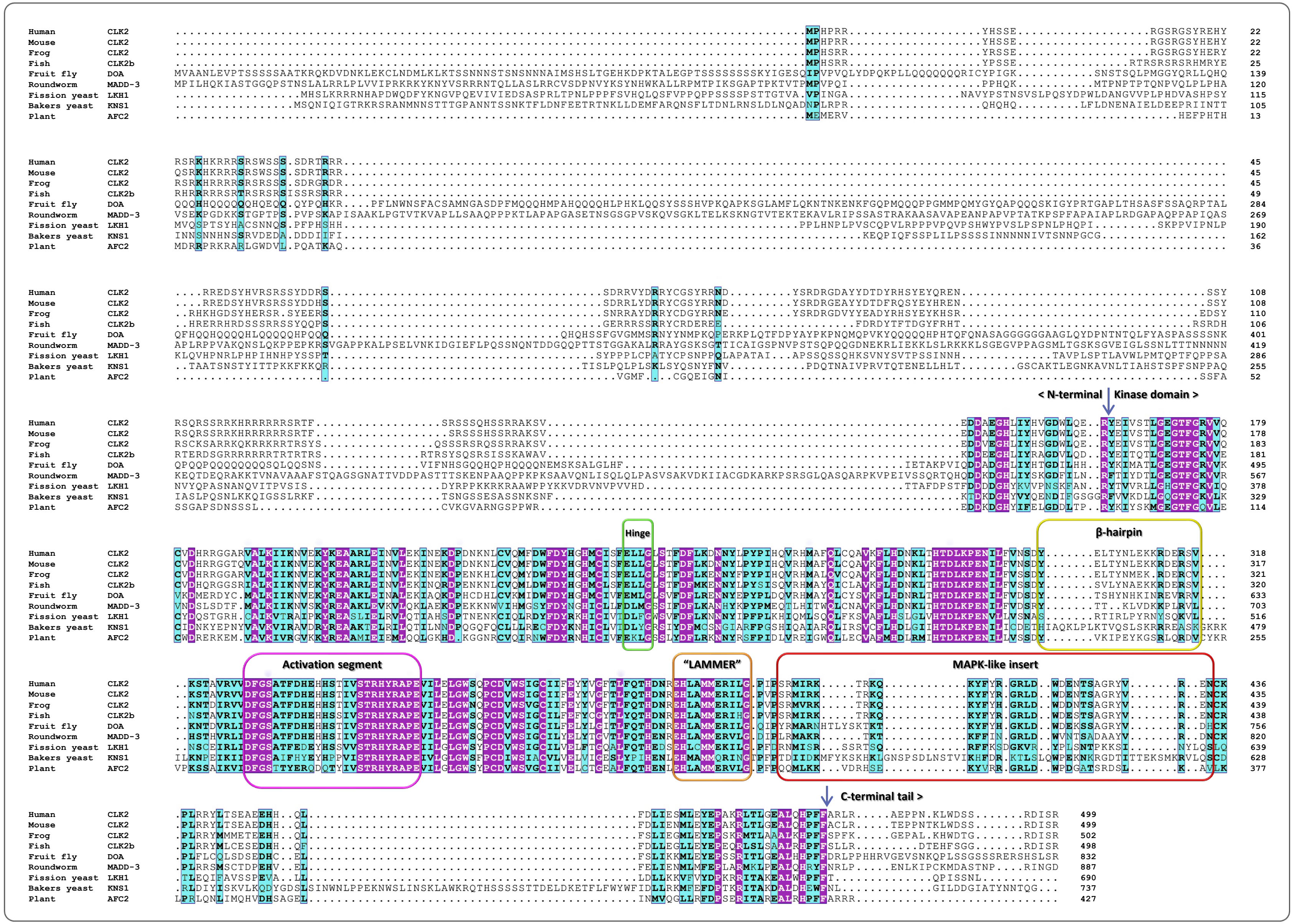
Human CLK2
Mouse CLK2
Frog CLK2
Fish CLK2b
Fruit fly DOA
Roundworm MADD-3
Fission yeast LKH1
Bakers yeast KNS1
Plant AFC2
< N-terminal
Kinase domain >
Hinge
β-hairpin
Activation segment
"LAMMER"
MAPK-like insert
C-terminal tail >

Figure S1.

| | |
|---|---|
| **CLK2**<br><br>Human | 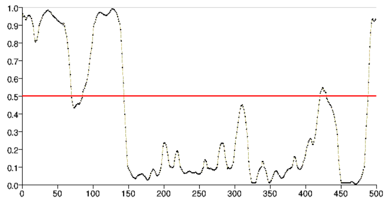 |
| **CLK2**<br><br>*Xenopus tropicalis*<br>(Frog) | 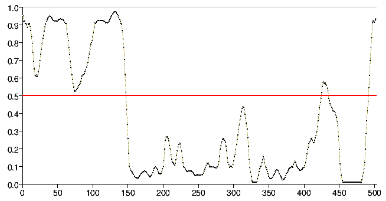 |
| **DOA**<br><br>*Drosophila melanogaster*<br>(Fruit fly) | 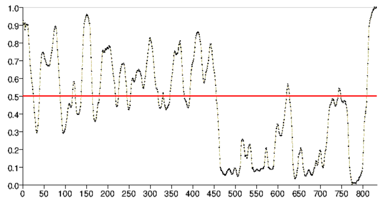 |
| **MADD-3**<br><br>*Caenorhabditis elegans*<br>(Roundworm) | 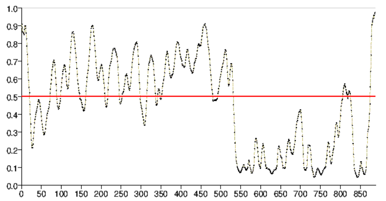 |
| **KNS1**<br><br>*Saccharomyces cerevisiae*<br>(Baker's yeast) | 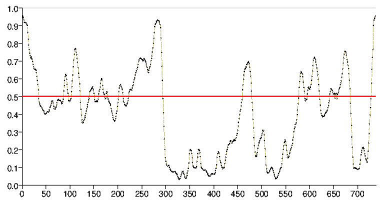 |
| **AFC2**<br><br>*Arabidopsis thaliana*<br>(Plant) | 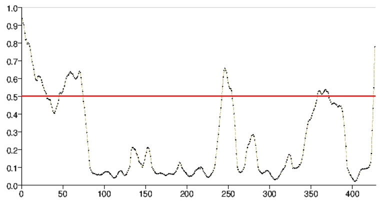 |

**Figure S2.**
