## Supplementary figures and images for "Comparison of the CDC2-like kinase family across eukaryotes highlights the functional conservation of these unique biological thermometers"

### Supplementary Figures 3

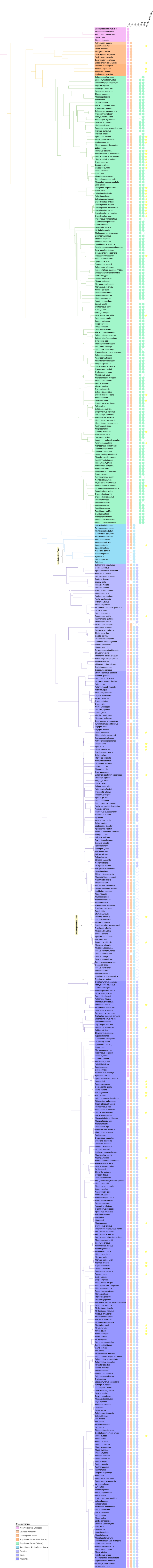
